# Individual differences in (dis)honesty are represented in the brain’s functional connectivity: Robust out-of-sample prediction of cheating behavior

**DOI:** 10.1101/2020.05.12.091116

**Authors:** Sebastian P.H. Speer, Ale Smidts, Maarten A.S. Boksem

## Abstract

Many of the economically most costly forms of unethical behavior such as tax evasion, stock manipulations or movie and music piracy relate to the moral domain of (dis)honesty, in which unethical behavior is not targeted at a clearly identifiable victim. While large individual differences in (dis)honesty are evident, the neurocognitive determinants of this heterogeneity remain elusive. We combined connectome-based predictive modelling (CPM) on resting state functional connectivity patterns with a novel experimental task, which measures spontaneous and voluntary cheating inconspicuously, to investigate how these task-independent neural patterns shape our (dis)honest choices. Our analyses revealed that functional connectivity in a network of regions, including the dorsolateral prefrontal cortex and the inferior frontal gyrus, commonly linked to cognitive control processes, but also the medial prefrontal cortex and temporal pole, associated with self-referential thinking, and the caudate nucleus, linked to reward processing, are of central importance in promoting honesty. In a leave-one-out cross-validation analysis, we show that this neural model can reliably and accurately predict how much an unseen participant will cheat on our task. Participants who cheated the most, also scored highest on several impulsivity measures, which highlights the ecological validity of our task. Notably, when comparing neural and self-report measures, the neural measures were found to be significantly more important in predicting cheating. Our findings suggest that a person’s dis(honest) decisions depend on how well the self-referential thinking network is functionally connected to the control and reward networks.

## Introduction

Cheating and dishonesty, manifested in diverse behaviors such as financial fraud, scientific misconduct and software piracy, is more prevalent than ever and represents one of the economically most costly forms of unethical behavior. However, it is evident that not everybody is a cheater: there are substantial individual differences in (dis)honesty, ranging from people who embody integrity and remain honest even when it comes at their own cost, such as Abraham ‘honest Abe’ Lincoln, to individuals such as Jordan ‘Wolf of Wallstreet’ Belfort, who greedily engaged in fraudulent stock market manipulations that led to investor losses of more than 200 million US Dollars.

In addition to such anecdotal evidence, several studies have shown that, when given the opportunity, individuals indeed differ considerably in how frequently they cheat (Gino et al., 2012; 2014, Speer, Smidts & Boksem, 2020). As in the example of the ‘Wolf of Wallstreet’, a greedy personality appears to be associated with dishonesty: research in social and personality psychology has found that greedy people indeed find a variety of moral transgressions more acceptable and engage in such unethical behaviors more frequently than less greedy people do (Seuntjens et al., 2019).

Yet, people do not only care about their own financial gains, which is evident from the omnipresence of prosocial behaviors such as altruism, reciprocity and honesty. When exposed to an opportunity to cheat, the way we view ourselves, our self-concept (Aronson 1969; Baumeister 1998; Bem 1972), may motivate us to refrain from cheating. People highly appreciate integrity and honesty in others and also have strong convictions of their own moral standards (Dhar & Wertenbroch, 2012). Violations of these moral standards would require a negative update of one’s self-concept and people are motivated to avoid this (Berthoz et al., 2006). As a result, people often tend to uphold their self-concept even if it means forgoing financial gains (Mazar, Amir, & Ariely, 2008).

The extent to which an individual focusses on their positive self-concept versus their greedy desires, and how this influences their moral decisions, may be associated with stable personality traits. For instance, the extent to which fairness concerns or social norms influence our judgement and decision-making may be linked to heterogeneity in (dis)honesty (Houser, Vetter & Winter, 2012; Cohn, Marechal & Noll, 2015; Cohn, Fehr & Marechal, 2014), while impulsivity and sensation-seeking have been associated with cheating (Anderman et al., 2009). In addition, it has been found that creative people cheat more (Gino et al., 2012, 2014), which may be driven by a greater creative ability to justify moral transgressions.

However, the neuro-cognitive processes underlying these individual differences have remained elusive. Insight into these mechanisms may provide us with a better and more detailed understanding of the processes responsible for the observed individual differences in dishonesty, and how they interact to shape our moral choices. A promising approach of investigating individual differences in (dis)honesty is to identify its neural correlates using resting state functional magnetic resonance imaging (rsfMRI). Since all individuals are unique, it is reasonable to expect that brain functional organization varies between individuals as well. Indeed, a study on a large dataset from the Human Connectome Project revealed that functional connectivity profiles can be used as a ‘fingerprint’ to identify individuals (Finn et al., 2015). Specifically, it has been shown that variability in whole brain functional connectivity is substantial across individuals, and that an individual’s functional connectome is robust and reliable across resting-state and task-based sessions over time and can even be reproduced between task and rest (Cao et al., 2014; Zuo and Xing, 2014; Finn et al., 2015). In previous research, rsfMRI has been employed to successfully link functional connectivity to individual differences in personality (Nostro et al., 2018; Cai et al., 2020) and social decision making and behavior, such as impulsivity in economic decision-making (Li et al., 2013), trust behavior (Hahn et al., 2014), reciprocity of a gift-giving (Caceda et al., 2015) and preference for social information (Zhang and Mo, 2016). In light of these findings, rsfMRI may provide added value to personality questionnaires as it provides direct access to the underlying cognitive and psychological sources driving this heterogeneity that is not distorted by social desirability bias (Grimm, 2010).

These underlying psychological and cognitive sources may not even be consciously accessible to the participants and may thus not be measurable with questionnaires.

Based on the great promise of resting state functional connectivity for investigating the underpinnings of individual differences in personality and decision making, this study examined whether the resting functional connectome can predict an individual’s propensity to cheat.

Recent neuroscientific evidence tends to support the notion of two opposing forces of greed and moral self-concept that steer us towards (dis)honesty. In a recent fMRI study (Speer, Smidts & Boksem, 2020), it was found that activity in the nucleus accumbens (Nacc), associated with reward anticipation and greed (Ballard & Knutson, 2009; Knutson, Adams, Fong, & Hommer, 2001; Abe & Greene, 2014), promotes cheating, particularly for individuals who tend to cheat a lot, whereas a network consisting of Posterior Cingulate Cortex (PCC), bilateral Temporoparietal Junctions (TPJ) and Medial Prefrontal Cortex (MPFC), associated with self-referential thinking processes (Gusnard et al, 2001; Meffert et al., 2013; Van Buuren et al., 2010), promotes honesty, particularly in individuals who are generally honest. In addition, numerous studies have proposed that cognitive control is needed to resolve this tension between reward and self-concept (Abe & Greene, 2014; Gino, Schweitzer, Mead, & Ariely, 2011; Greene & Paxton, 2009; Maréchal, Cohn, Ugazio, & Ruff, 2017; Mead, Baumeister, Gino, Schweitzer, & Ariely, 2009). In accordance with these findings, the study by Speer et al. (2020) revealed that activity in cognitive control regions, namely the anterior cingulate cortex (ACC) and the inferior frontal gyrus (IFG; Swick et al., 2008; Carter & Van Veen, 2007) were recruited to resolve the conflict between self-interest and self-image.

In light of this previous research, we hypothesize that higher functional connectivity within the self-referential thinking network, including the MPFC, TPJ, PCC and temporal poles will be predictive of honesty. Similarly, we expect that higher connectivity between cognitive control regions, such as the dlPFC, ACC or IFG on the one hand and the self-referential thinking network on the other hand will be additionally predictive of honesty, by further enhancing the influence of our self-concept on moral decisions. Conversely, higher connectivity within reward regions, such as the Nacc, caudate nucleus and ventromedial prefrontal cortex (vmPFC), is hypothesized to predict dishonesty, while a negative coupling between cognitive control and reward networks may be additionally predictive of dishonesty.

To test these hypotheses, we used a sample of 99 participants who completed a resting state scan and the ‘Spot-The-Difference Task’ (see Gai & Puntoni, 2017; Speer, Smidts & Boksem, 2020). This is an innovative task in which participants could cheat repeatedly, deliberately and voluntarily without suspicion of the real purpose of the task. We employed connectome-based predictive modeling (CPM) to investigate whether (dis)honesty can be reliably predicted from an individual’s unique pattern of functional connectivity. CPM has recently been developed to predict individual differences in human behavior (e.g., cognitive abilities & personality traits) from patterns of whole-brain functional connectivity (Finn et al., 2015; Rosenberg et al., 2016; Shen et al., 2017). Specifically, the predictive power of CPM has been illustrated in research on fluid intelligence (Finn et al., 2015), sustained attention (Rosenberg et al., 2016), and creativity (Beaty et al., 2018) revealing reliable prediction of these behavioral variables in participants whose data were not used in model fitting. Importantly, this approach differs from methods implemented in previous rsfMRI studies as it uses cross-validation to assess predictive accuracy instead of just establishing correlational relationships. As the standard CPM approach does not allow to identify the importance of each of the individual connections, we extended the CPM by integrating multiple regression and lasso-regression. This allowed us to determine which individual connections are the most important ones in predicting an individual’s propensity to cheat. Moreover, this extended CPM approach, combined with permutation importance analysis, enabled us to directly compare the importance of personality questionnaire measures with functional connectivity measures in predicting cheating propensity.

## Methods

### Participants

The reported analyses are based on 99 participants (67 females; 24 nationalities; age 18-44 years; *M* = 24.3, *SD* = 4.26) from three separate studies. The reason for running three studies was driven by our motivation to obtain a large and diverse sample size. The first sample of participants consisted of a student sample (N=49, 37 females; age 18-35 years; *M* = 24.4, *SD* = 3.34) from now on referred to as Study 1. The second sample was collected from a student sample at another university in a different scanner (N=9, 8 females; age 21-24 years; *M* = 21.8, *SD* = 1.12). The third sample (Study 3) consisted of a general population sample from a different city recruited via flyers around town and online via Facebook (N=41, 23 females; age 18-43 years; *M* = 24.8, *SD* = 5.4) and resting state data was collected in a different scanner. No significant differences in demographics (age and gender) were identified between the samples. All participants were right-handed with normal or corrected to normal vision, and no record of neurological or psychiatric diseases. The studies were approved by the respective university Ethics Committees and were conducted according to the Declaration of Helsinki.

### Task and Stimuli

#### Spot-The-Difference Task

As described in a previous study by Speer, Smidts and Boksem (2020), in the Spot-The-Difference task, participants were presented with pairs of images and were instructed that there were always three differences present between the image pairs. Differences consisted of objects that were added to or removed from an image, or objects that differed in color between images. However, images could actually contain one, two, or three differences. Participants were requested to find three differences between the images. Since reward (see below) was contingent on participants *reporting* that they had found all three differences, without having to point them out, this design allowed and encouraged cheating behavior (i.e., reporting having found all three, even when objectively fewer than three differences were present in the images).

Participants were instructed that the purpose of the study was to investigate the underlying neural mechanisms of visual search for marketing purposes such as searching for a product in an assortment or information on a webpage. In order to increase credibility of this cover story a simple visual search task was added at the beginning of the experiment (see Appendix 1). Further, participants were instructed that the neurocognitive effect of motivation, elicited by monetary reward, on speed and accuracy of visual search would be investigated. Although participants were told that there were three differences in all trials, in 25% of the trials there were only two differences and in 25% there was only one difference. All stimuli were standardized in size and were presented on a white background on a computer screen. The ratio of 50% −50% (three differences vs less than three differences) was chosen based on the results of pilot studies that indicated this ratio to be optimal in reducing suspicion that the pairs did not always contain three differences.

Trials were further categorized into normal (50%), hard (25%) and very hard trials (25%), for which participants could receive 5cts, 20cts, and 40cts, respectively. All the trials with three differences (the filler trials) were categorized as normal trials, whereas trials with less than three differences (the trials of interest) were randomly categorized as hard or very hard trials. Consequently, the reward was independent of the number of differences in the image pair for the trials of interest, which is important in order to be able to disentangle the effects of reward and cheating magnitude (the actual number of differences) on cheating behavior. The different levels of difficulty were added to reduce suspicion about the real purpose of task. It was assumed that if trials are labeled as hard or very hard, it would be more credible to the participant that the image pair actually contained three differences, but they were just too hard to spot. In addition, levels of difficulty were introduced to eliminate possible demand effects: we wanted participants to cheat for monetary reward and not to prevent seeming incompetent, which may be associated with different underlying neural mechanisms and consequently confound the analysis.

To further reduce suspicion about the purpose of the study, approximately 10% of all trials were point-and-click trials. In these trials, participants had to click on the location in the images where they spotted the differences using a joy-stick. Consequently, cheating was not possible on the point-and-click trials. Participants always knew prior to the start of a trial whether it was a point-and-click trial indicated by a screen requesting participants to click on the image. This ensured that participants would not refrain from cheating on all other trials, while still reducing the suspicion about the real purpose of the study. Participants were told that only 10% of trials were point-and-click trials because it would take too much time to point out the differences for every pair. In sum, there were 144 regular trials (of which 72 cheatable trials) and 12 point-and-click trials. The maximum amount of money earned, in case a participant cheated on all cheatable trials was approximately 35 Euros, whereas in case a participant would not cheat at all he or she would earn approximately 7.50 Euros. To be fair to all participants, after completion of the full study, participants were debriefed and they were all paid out the maximum amount, irrespective of their actual cheating behavior.

Each trial started with a fixation cross which was presented for a variable amount of time between 1-3s (see Figure 1). Subsequently, the Level of Difficulty screen was presented for 2 seconds informing the participants about the level of difficulty of the upcoming trial. This screen also displayed how much money could be earned on that trial. As a result, participants were constantly aware of the potential gains of cheating. Next, an image pair was presented for 6s, a length determined by the behavioral pilots, and participants engaged in the visual search. Afterwards, the participants were asked whether they spotted all three differences (yes/no response). On this decision phase screen, again the potential reward for this trial was presented, in order to make the reward more salient and increase cheating behavior. After 3s, the response phase started in which participants’ responses were recorded. In the decision phase and the response phase the current balance was also shown, which was done to demonstrate to the participants that if they stated that they had found the three differences, their current balance increased immediately. It was assumed that this direct noticeable effect of behavior on the increase of the current balance, would further motivate participants to cheat.

**Figure 1.**
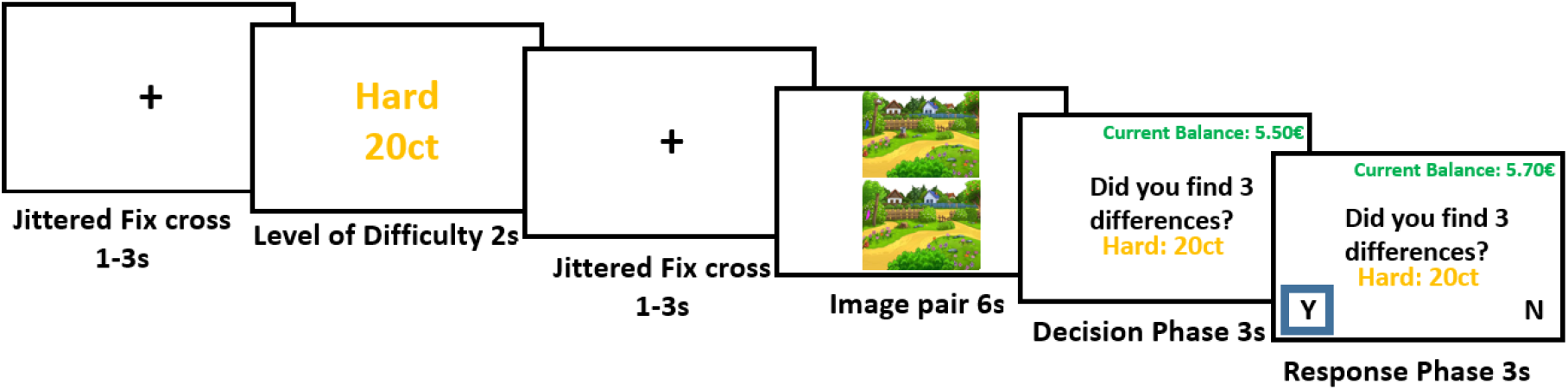
One trial of the Spot-The-Differences paradigm. Participants viewed a screen indicating the difficulty and value of the trial, then the image pair appeared for six seconds and then participants had to indicate whether or not they spotted all three differences.

The buttons corresponding to “Yes” and “No” were switched across trials to reduce the response bias for the dominant hand. Once the participants responded, the choice was highlighted by a blue box for 500ms to indicate that the response was recorded, and the trial ended. If no response was made, the trial ended after 3s. In addition, there were five practice trials, in which participants could get acquainted with the task. Stimulus presentation and data acquisition was performed using Presentation® software (Version 18.0, Neurobehavioral Systems, Inc., Berkeley, CA, www.neurobs.com).

#### Stimuli

Stimuli for the task consisted of 144 Spot-The-Difference image pairs that were downloaded from the Internet. Cartoon images of landscapes containing several objects were selected, to make them engaging and challenging enough for the participants. Landscapes were chosen as they generally satisfied the necessary criterion of containing several different objects. The stimuli consist of pairs of images that are identical apart from a certain number (1-3) of differences that were created using Adobe Photoshop. Differences consisted of objects added to or removed from the landscape picture or changed colors of objects. Differences were fully randomized across all pairs of images, which means that all image pairs could be presented with either one, two or three differences. To make sure that participants would be able to find the differences between the images in a reasonable amount of time, we ran a pilot study on Amazon’s Mechanical Turk (N=205) to test the difficulty to spot the differences between the images and to determine the optimal duration of picture presentation (see Appendix 2).

#### Experimental procedure

All participants were first informed about and checked on the safety requirements for MRI scanning. They then completed the resting state scan. Before the Spot-The-Difference task started, participants were then introduced to the cover story, and the tasks and they signed the informed consent form. Subsequently, they completed practice trials for both visual search tasks. Afterwards, the participants completed the simple visual search task (5 min) followed by the Spot-The-Difference task which took approximately 40 minutes. After completing the two tasks participants filled-in a short questionnaire including questions about their thoughts on the purpose of the task.

After completion of the experimental session, participants received an email with a link to a Qualtrics questionnaire including measures for impulsivity, greed, creativity, manipulativeness and sensitivity to different moral foundations (explained below), which they were allowed to fill out at home. First, in the test battery, we included four impulsivity scales: a) the Brief Sensation Seeking Scale (BSSS; Hoyle, Stephenson, Palmgreen, Lorch & Donohew, 2002), b) the BisBas scale to assess dispositional inhibition and approach behavior (Carver & White, 1994), c) the short version of the UPPS-P Impulsive Behavior scale (Cyders, Littlefield, Coffey, & Karyadi, 2014) and d) a Risk seeking scale implementing a standard risk preference elicitation method where one can choose between a certain amount of money and a risky gamble. To quantify risk preference, participants were presented with sequence of binary choices between a certain amount of money for sore or a gamble with a fifty percent chance of winning 30€ for sure and a fifty percent of not winning anything. Whereas the gamble remains the same for each question the amount gained for sure increases at each step. A person’s risk preference could thus be established by identifying the amount of money for sure at which the person switches from the gamble to certain payout (example item: “Would you prefer 13€ for sure or 0€ or 30€ with a 50-50% chance”). These impulsivity scales were selected because dishonesty and cheating has been linked to impulsivity as a personality trait (Anderman et al., 2009).

Second, we measured how an individual’s sensitivity to different moral foundations, namely Care vs. Harm, Fairness vs. Cheating, Loyalty vs. Betrayal, Authority vs. Subversion, Sanctity vs. Degradation and Liberty vs. Oppression, may influence cheating by including two such measures: a) the Moral Foundations Questionnaire (MFQ; Graham et al., 2011) and b) the Moral Foundations Vignettes (Clifford, Iyengar, Cabeza & Sinnott-Armstrong, 2015).

Third, as greed is assumed to drive cheating behavior (Seuntjens et al., 2019), the Dispositional Greed Scale was added (Seuntjens et al., 2015). Fourth, an individual’s creativity was measured by means of three scales: a) the Remote Associates Test (Mednick, 1968), b) Gough’s Creative Personality Scale (gough, 1979), and c) Hovecar’s Creative Behavior Inventory (Hovecar, 1979), as it has been found that more creative people tend to cheat more (Gino et al., 2012). Fifth, the MACH-IV test (Christie & Geis, 1970) to measure manipulativeness was added as Machiavellianism has also often been associated with unethical behavior (Tang & Chen, 2008). Participants were informed that they would only receive their payment once they completed the questionnaires.

### FMRI acquisition

For Study 1, the functional magnetic resonance images were collected using a 3T Siemens Verio MRI system. Functional scans were acquired by a T2*-weighted gradient-echo, echo-planar pulse sequence in descending interleaved order (3.0 mm slice thickness, 3.0 × 3.0 mm in-plane resolution, 64 × 64 voxels per slice, flip angle = 75°). TE was 30ms and TR was 2030ms. A T1-weighted image was acquired for anatomical reference (1.0 × 0.5 × 0.5 mm resolution, 192 sagittal slices, flip angle = 9°, TE = 2.26ms, TR = 1900ms).

For Study 2, the functional magnetic resonance images were collected using a 3T MRI system (General Electric). Functional scans were acquired by a T2*-weighted gradient-echo, echo-planar pulse sequence in ascending interleaved order (3mm slice thickness, 3.5 mm slice gap, 3 × 3 mm in-plane resolution, 64 × 64 voxels per slice, flip angle = 75°, TE = 30ms, TR = 2030ms). A T1-weighted image was acquired for anatomical reference (1.0 × 1.0 × 1.0 mm resolution, 160 sagittal slices, TE = 2.35ms, TR = 7.21ms).

For Study 3, the functional magnetic resonance images were collected using a 3T Phillips Achieva MRI system. Functional scans were acquired by a T2*-weighted gradient-echo, echo-planar pulse sequence in descending interleaved order (3.0 mm slice thickness, 3.0 × 3.0 mm in-plane resolution, 64 × 64 voxels per slice, flip angle = 76°). TE was 27ms and TR was 2000ms. A T1-weighted scan was acquired using 3D fast field echo (TR: 82◻ms, TE: 38◻ms, flip angle: 8°, FOV: 240◻×◻188◻mm, in-plane resolution 240 × 188, 220 slices acquired using single-shot ascending slice order and a voxel size of 1.0◻×◻1.0◻×◻1.0◻mm).

For all studies, the stimuli were presented using Presentation® software (Version 18.0, Neurobehavioral Systems, Inc., Berkeley, CA, www.neurobs.com).

### Preprocessing

Data was preprocessed using the standard pipeline of the CONN toolbox (https://www.nitrc.org/projects/conn) in MATLAB. This pipeline includes realignment of the functional data using SPM12’s realign & unwarp procedure (Anderson et al., 2001), where all scans are coregistered and resampled to a reference image (first scan of the first session) using b-spline interpolation. Subsequently, outlier detection was performed from the observed global BOLD signal and the amount of subject-motion in the scanner. Acquisitions with framewise displacement above 0.9mm or global BOLD signal changes above 5 SD are marked as potential outliers. Framewise displacement is computed at each timepoint by considering a 140×180×115mm bounding box around the brain and estimating the largest displacement among six control points placed at the center of this bounding-box faces. Afterwards, both the functional and the anatomical data are normalized into standard MNI space and segmented into grey matter, white matter, and CSF tissue classes using SPM12’s unified segmentation and normalization procedure (Ashburner and Friston, 2005). As a last step, functional data is smoothed using spatial convolution with a Gaussian kernel of 8mm full width half maximum (FWHM), in order to increase BOLD signal-to-noise ratio and reduce the influence of residual variability in functional and gyral anatomy across subjects.

As a next step denoising of the functional data was performed again using the standard pipeline from the CONN toolbox. For each participant, CONN implemented CompCor, a method for identifying principal components associated with segmented white matter (WM) and cerebrospinal fluid (CSF). In a first-level analysis, aCompCor components (Behzadi et al., 2007) and first-order derivatives of motion were entered as confounds and regressed from the BOLD signal. In addition, preprocessing steps included temporal band-pass filtering (0.008 Hz – 0.09 Hz), linear detrending, and regression of outlying functional volumes (>97th percentile in normative sample; global-signal z-value threshold = 5, subject-motion mm threshold = 0.09) identified using the artifact removal toolbox (ART) (https://www.nitrc.org/projects/artifact_detect/).

#### Functional Network Construction

To define brain regions of interest we used dictionary learning (Mensch et al., 2016, Mensch et al., 2018) to extract 80 components from the denoised resting state data. Dictionary learning is a sparsity-based decomposition method for extracting spatial maps. It extracts maps that are naturally sparse and usually cleaner than ICA (Mensch et al., 2016), and it has been found to be the method that leads to the highest predictive success in a comparison of different connectome-based prediction pipelines (Dadi et al., 2019). In addition, the authors found that 80 components are the optimal amount for predictive performance. Subsequently, a Random-walk based extraction of regions from the brain networks obtained by the dictionary learning algorithm was used as proposed in Abraham et al. (2014) resulting in 142 regions. To estimate functional connectivity between these 142 regions efficiently, we use the Ledoit-Wolf regularized shrinkage estimator (Ledoit and Wolf, 2004; Varoquaux and Craddock, 2013), which gives a closed form expression for the shrinkage parameter. For parametrization of the functional interactions, Pearson’s correlation was used.

**Figure 2.**
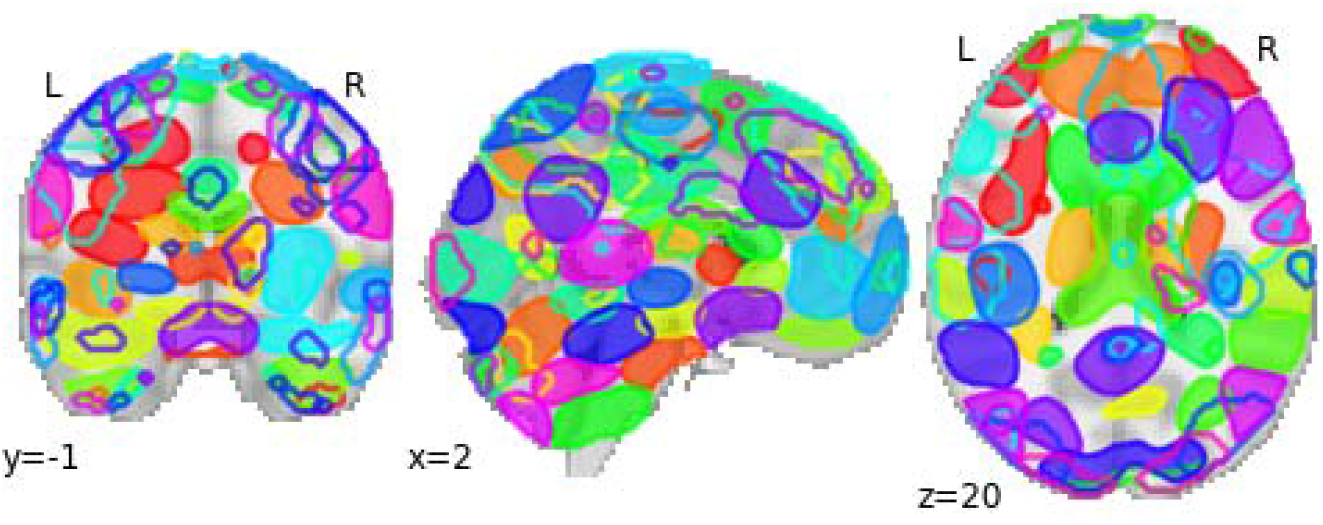
A total of 80 components extracted from the dictionary learning algorithm

#### Connectome-Based Predictive Modeling

The main analysis utilized CPM to predict participants’ propensity to cheat from whole-brain resting state functional connectivity patterns. CPM is a recently developed tool for identifying functional brain connections related to a behavioral variable of interest, which are then used to predict behavior in unseen participants (i.e., participants whose data were not used in the training of the model; Shen et al., 2017). CPM was introduced and described in recent neuroimaging literature in a series of studies reporting its successful implementation in prediction of cognitive variables such as fluid intelligence, attention control and creativity (Finn et al., 2015; Rosenberg et al., 2016; Shen et al., 2017, Beaty et al., 2018). The MATLAB syntax used for CPM is freely available online (https://www.nitrc.org/projects/bioimagesuite/).

We implemented the CPM procedure in Python since the functional network construction steps were also performed in Python with use of the Nilearn package (Abraham et al., 2014b). We extended the standard procedure, described in detail below. As in the standard CPM approach (Shen et al. 2017), as a first step, the level of honesty which is the reverse of the number of times a participant cheated on the Spot-The-Difference task (the cheatcount), was correlated with each edge (i.e., correlation of mean BOLD signals between a given pair of brain regions) in the functional connectivity matrix of each participant (see Figure 3A&B). Subsequently, a threshold was applied to the connectivity matrix to keep only the edges that were significantly positively or negatively correlated with honesty (p<0.001, see Figure 3C). The positive edges, from now on referred to as the honesty network (as these edges are positively correlated with honesty) and the negative edges, referred to as the dishonesty network (as these are negatively correlated with honesty), will from this stage onwards be analyzed separately.

**Figure 3.**
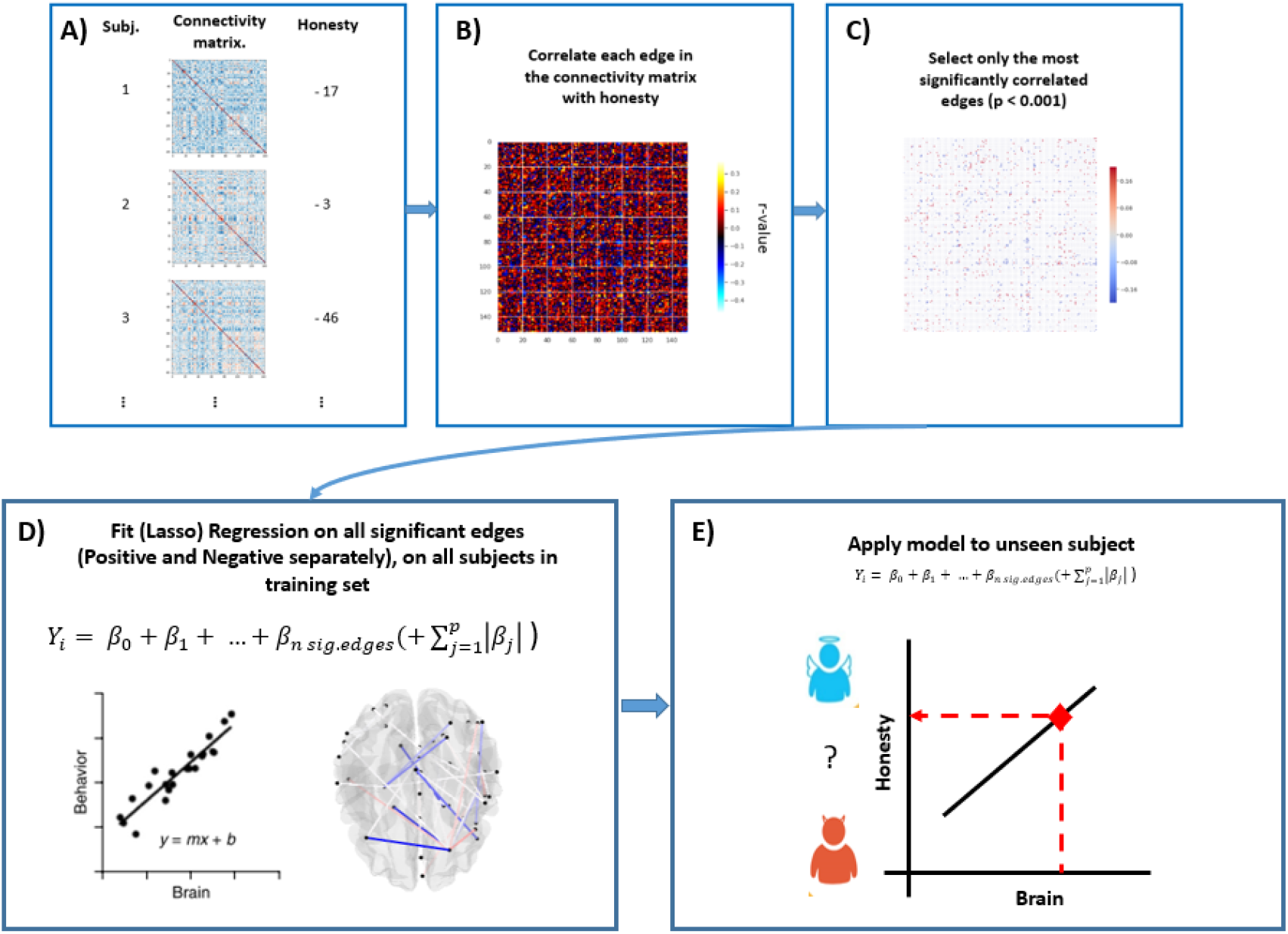
Adapted CPM procedure: A-B) As a first step, the correlation between each edge in the connectivity matrix and level of honesty is computed. C) Next, the correlations are thresholded and only the significant edges are retained. D) We then fit a (lasso) regression model using the selected edges to the level of honesty on the participants in the training set. E) As a last step, we use the model to predict the level of honesty of the left-out subject (whether someone is more a saint = honest participant or a devil = person who cheats a lot/ cheater).

In our extended procedure, we then fit a multiple regression model and a lasso regression model on the significant edges, to derive beta coefficients for each of the edges (see Figure 3D). This allows us to assess the importance of the different edges that are entering the predictive model. Afterwards, a linear regression model was specified to estimate the relationship between the cheatcount predicted by the model and the observed cheatcount. Lastly, as in the standard approach, the model was applied to unseen participants in a leave-one-out cross validation scheme (see Figure 3E). Specifically, all steps described, including correlating edges with honesty and feature selection (thresholding) and model estimation, were implemented on n – 1 participants connectivity matrices and cheatcount scores, and then the fitted model was tested on the left-out participant.

Due to the fact that feature selection is performed inside the cross-validation loop, slightly different edges may be selected at each iteration which result in slight variations in the predictive models. The predictive power of the model is assessed by means of the statistical significance of the Pearson correlation between the predicted honesty scores and the observed scores. Statistical significance of this correlation is estimated by means of permutation testing where honesty scores are permuted and the CPM procedure is repeated 1000 times. The empirical score is then compared against the null distribution to derive a p-value.

To estimate feature importance for the multiple regression and Lasso models, a permutation-importance approach was implemented (Breiman, 2001). Specifically, in each cross validation iteration of the CPM procedure, the estimation of the predictive model was repeated and each of the predictors (edges) were permuted 5 times in sequence and the average prediction across the 100 permutations was recorded for each of the predictors. Finally, after CPM was performed, only the predictors that occurred in each prediction fold were selected and the reduction in correlation associated with permuting each coefficient (as compared to the correlation including the unpermuted predicted) at each fold was computed.

Feature importance was then calculated as the difference in correlation between predicted and actual level of honesty between the baseline model (no permuted predictors) and the permuted model. Consequently, the higher the difference, the more important the predictor was.

## Results

### Behavioral results

Substantial individual differences in the total amount of cheating were observed (Mean= 37%, Median=28%, SD=32%; see Figure 4): some participants cheated only on one or two trials (11% of participants), whereas others only missed one or two opportunities to cheat (4%).

**Figure 4.**
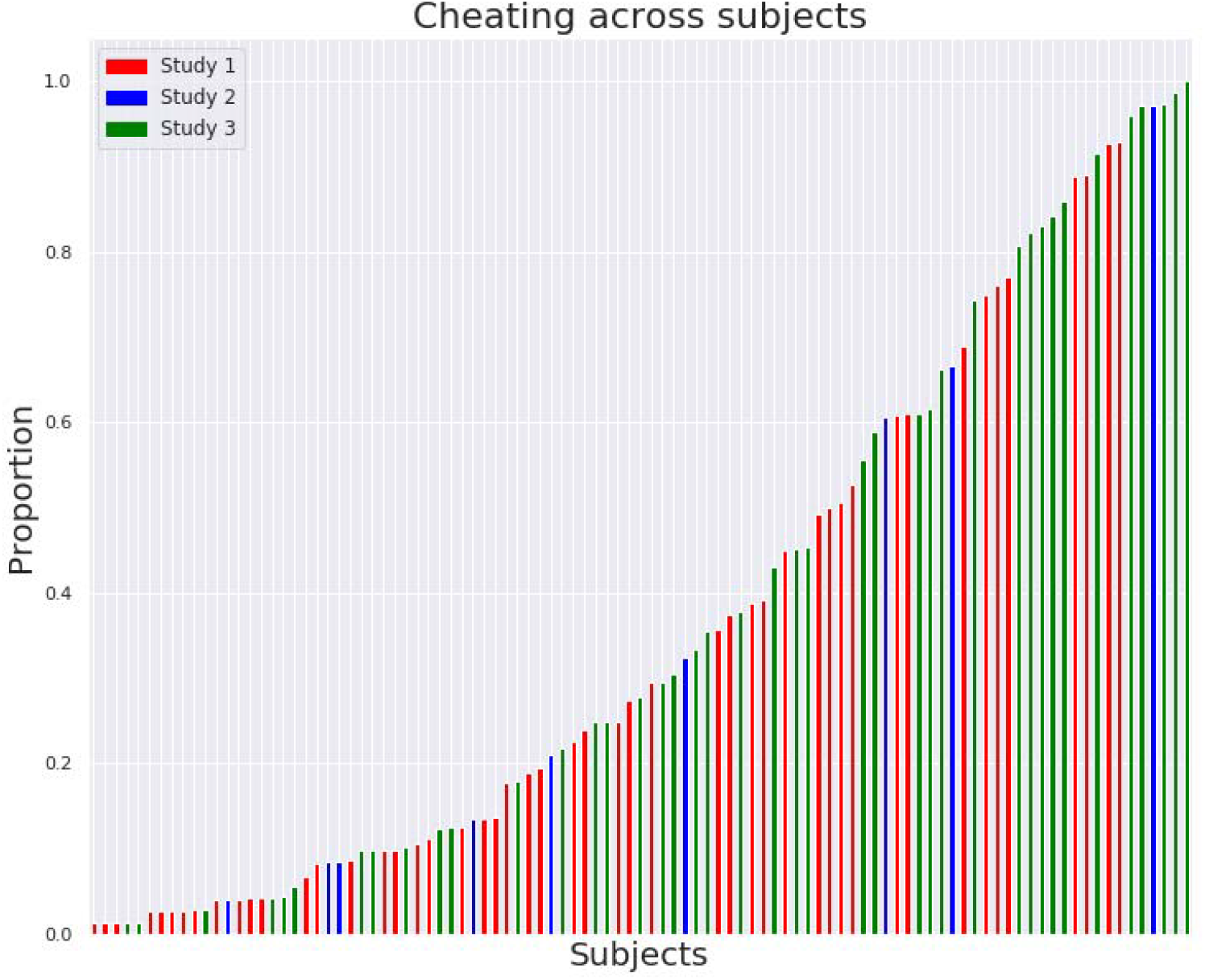
Individual differences in proportion of cheating on the Spot-The-Difference task. N= 99.

To assess suspicion about the real purpose of the study, participants were asked what the goal of the experiment was. Participants mentioned marketing research, consumer decision-making and visual search as our general cover story suggested that visual search is important for quickly locating one’s favorite brand or product in a supermarket. However, 12 participants mentioned dishonesty, moral decision making or related concepts, which suggests that they were suspicious of the real goal of the study. Importantly, no significant differences in cheatcount were found between suspicious and unsuspicious participants (*t* = 0.11, *p*=0.91), which suggests that suspicion about the purpose of the study did not have significant effects on cheating behavior. To assess the robustness of our findings, the neural analyses were conducted with all participants included and with the suspicious participants removed. The robustness checks revealed that CPM predictions remain significant also without suspicious participants (see Appendix 3).

We tested whether the magnitude of (dis)honesty we measure with our task generalizes to stable personality characteristics, to explore the ecological validity of the Spot-The-Difference task. To test this, we correlated the individual differences in (dis)honesty with scores from measures of impulsivity, moral foundations, greed, creativity and manipulativeness, respectively. We found that levels of honesty (reverse cheatcount) negatively correlated (for all correlations see Appendix 4) with four measures of impulsivity: a) The Brief Sensation Seeking Scale (r = −0.23, p < 0.05; Example item: “I would like to try bungee jumping”; Hoyle, Stephenson, Palmgreen, Lorch & Donohew, 2002), b) the positive urgency subscale of the short version of the UPPS-P Impulsive Behavior scale (r = −0.24, p < 0.05; Example item: “I tend to act without thinking when I am really excited.”), c) the lack of premeditation subscale (r = −0.21, p < 0.05; Example item: “I like to stop and think things over before I do them.”, reverse coded) of the short version of the UPPS-P Impulsive Behavior scale (Cyders, Littlefield, Coffey, & Karyadi, 2014), and d) the Risk seeking scale (r = −0.25, p < 0.05; Example item: “Would you prefer 13€ for sure or 0€ or 30€ with a 50-50% chance”). In accordance with previous literature (Anderman et al., 2009), these findings suggest that cheating is associated with higher impulsivity.

Unexpectedly, we also found that more honest participants scored higher on reward responsiveness of the BisBas scale than more dishonest participants (r = 0.21, p < 0.05; Example item: “When I get something I want, I feel excited and energized.”). In addition to impulsivity, and perhaps surprisingly, honesty also correlated negatively with how sensitive participants reported themselves to be to social norms (r = −0.32, p < 0.05; Example item: “You see a woman answering a phone call with the word “goodbye” instead of “hello”, 5 point response scale: not at all wrong – extremely wrong; Clifford, Iyengar, Cabeza & Sinnott-Armstrong, 2015), and to violations of purity (r = −0.25, p < 0.05; Example item: “Chastity is an important and valuable virtue”).

### Predicting (dis)honesty using a resting state functional connectome

As explained in the methods section, the CPM procedure identifies functional connections that are significantly positively related to honesty, which we term the honesty network, and edges that are significantly negatively related to honesty, which we term the dishonesty network. To identify the most important edges within the networks we used the multiple regression CPM and found a strong correlation for the honesty network between predicted and actual honesty (*r* = 0.61, *p_perm_*<0.001, see Figure 5C). The higher the functional connectivity within the honesty network, consisting of the dorsolateral prefrontal cortex (dlPFC), the medial prefrontal cortex (MPFC), the inferior frontal gyrus (IFG), the supplemental motor area (SMA), the temporal pole, the posterior cingulate cortex (PCC) and the caudate nucleus, the more honest the participants were (see Figure 5). No significant correlations were found for the dishonesty network. Consequently, we will from now on focus on how well the functional connections predict honesty (reverse cheatcount).

**Figure 5.**
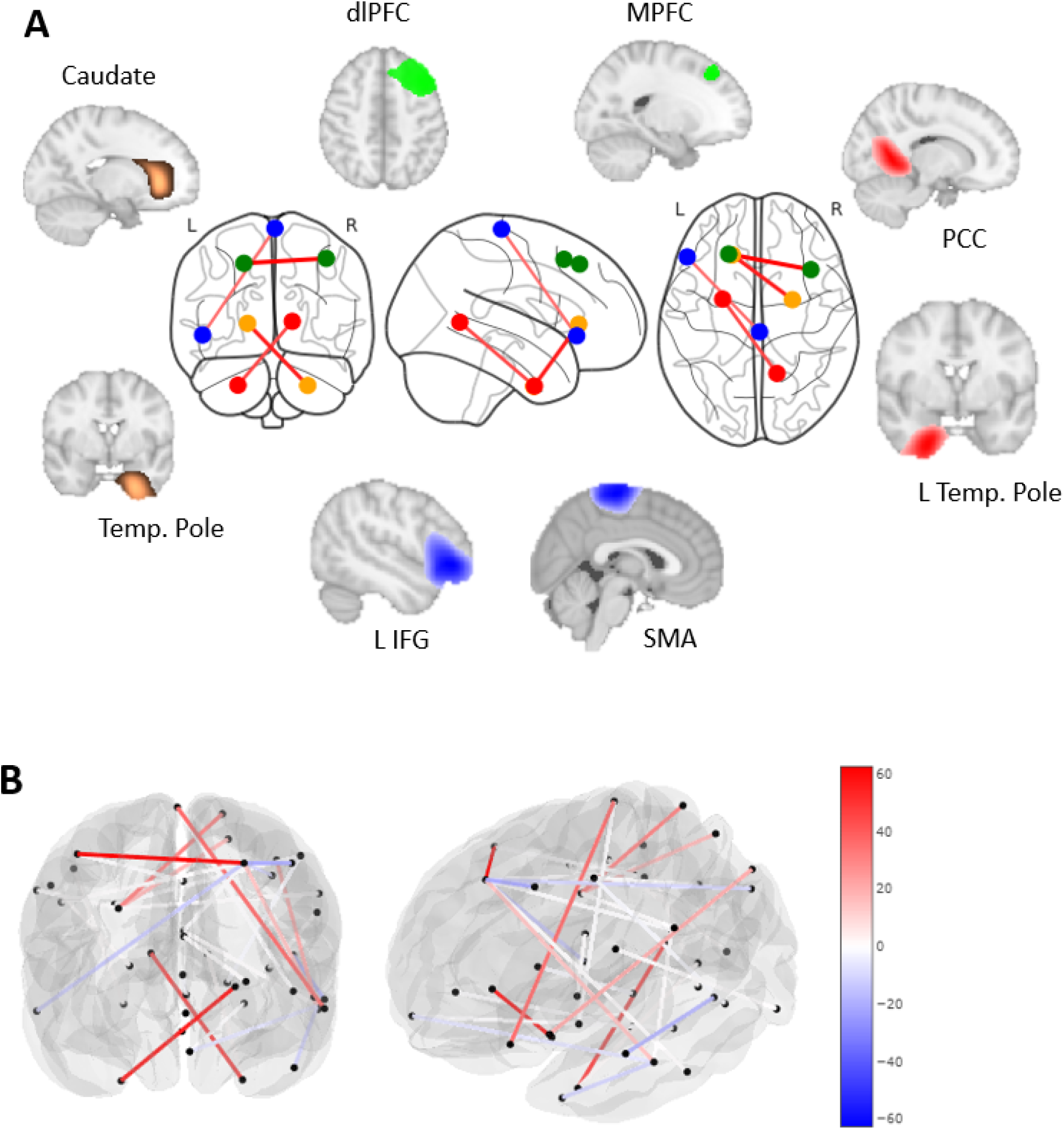
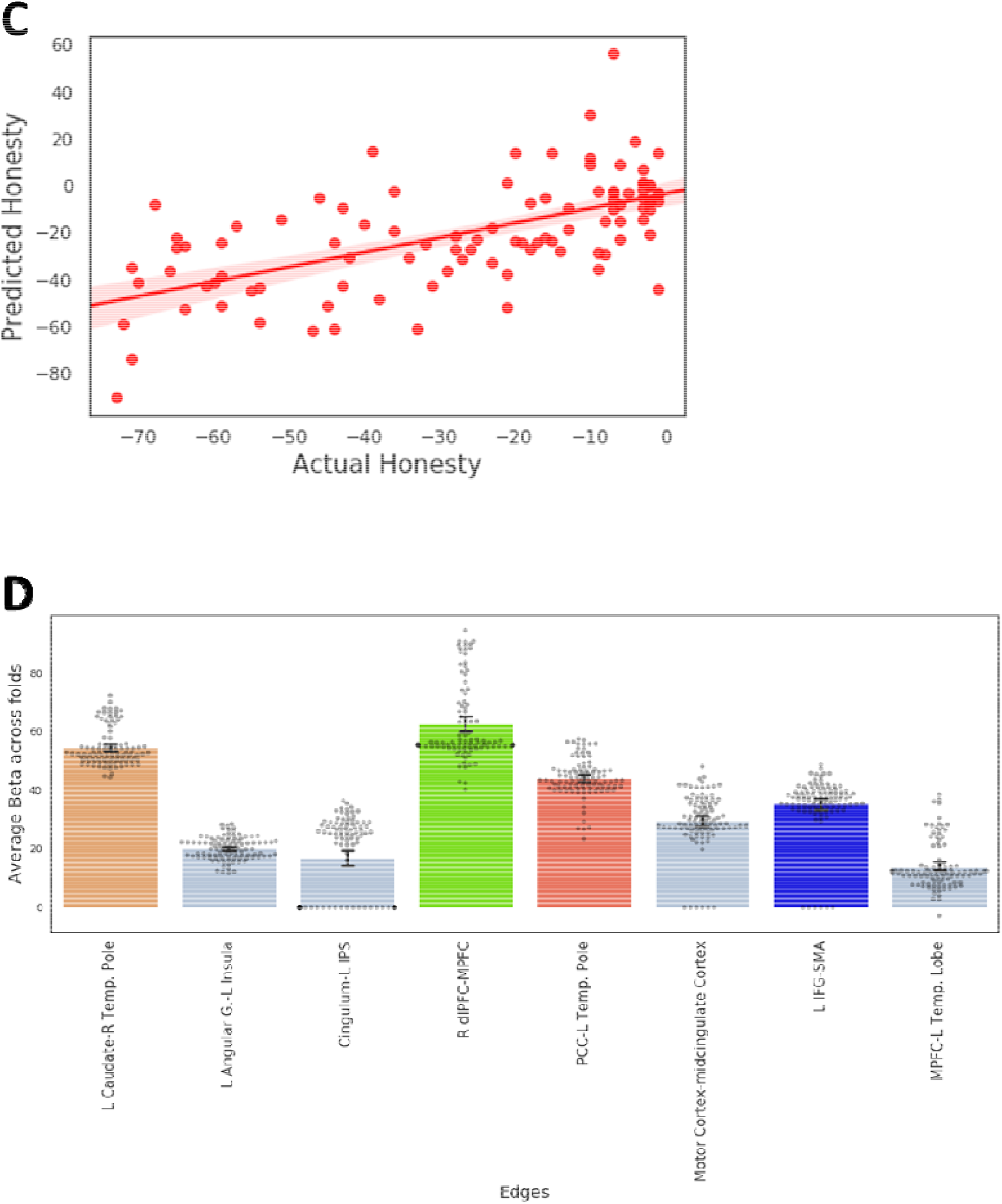
Higher functional connectivity in the honesty network is linked to more honest responses (lower cheatcount) in the Spot-The-Difference task. A) Connectome of the 4 strongest predictors (edges) of honesty (negative predictor of cheatcount) averaged across folds including connectivity between the left caudate and right Temporal Pole, the right dlPFC and MPFC, the PCC and left Temporal Pole and the left IFG and the SMA. B) Showing the full connectome including all edges selected during CPM. C) Correlation between predicted cheatcount and actual cheatcount. D) Barplot of beta coefficients for all edges of the honesty network averaged across folds that were significantly different from zero (colors correspond with Plot A; new edges depicted in grey, individual points reflect beta at each fold).

### Comparing predictive importance of the questionnaires to the functional connectome

As a next step, we investigated whether the neural data or the questionnaire data were better predictors of honesty in unseen participants. First, we had to establish whether successful prediction of honesty in unseen participants was possible using just questionnaire data. Since some participants did not complete all questions in the questionnaires, only 91 participants were included in this analysis. Similar to the connectome analysis, to identify the most important personality measures in predicting out-of-sample honesty, we employed the multiple regression CPM, which also led to a significant prediction accuracy (*r* = 0.42, *p*<0.05). However, the prediction accuracy was significantly lower than for the neural predictors (*z*= −1.77, *p*<0.05). We found that particularly self-reported impulsivity, namely our Risk Seeking scale (standard risk preference elicitation method) and the Positive-Urgency-subscale of the short UPPS Impulsive behavior scale, are on average (across folds) significant negative predictors of honesty (high cheatcount, see Figure 6A). In addition, an individual’s sensitivity to social norms was observed to be an important, although perhaps unexpected, negative predictor of honesty (see Figure 6A).

**Figure 6.**
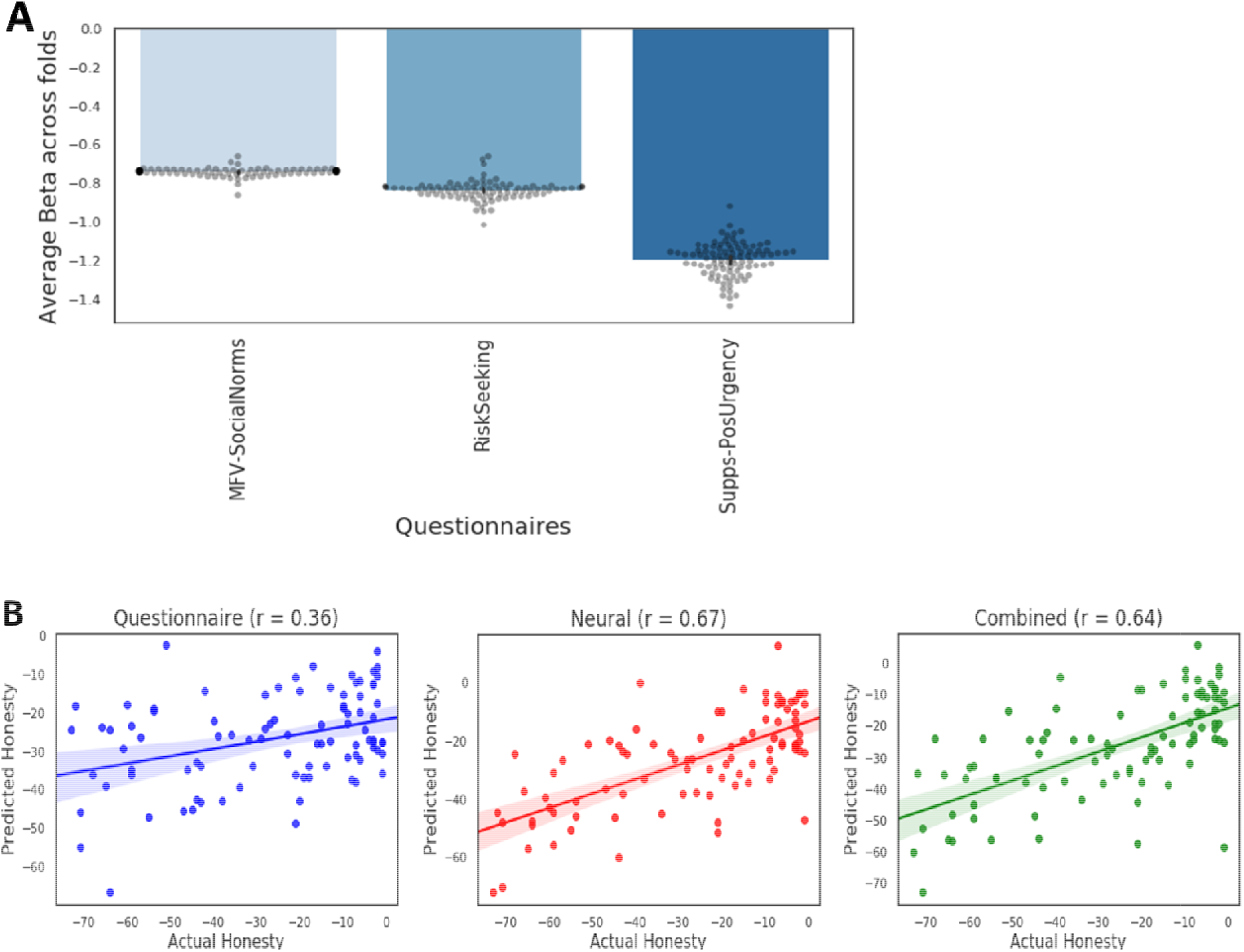
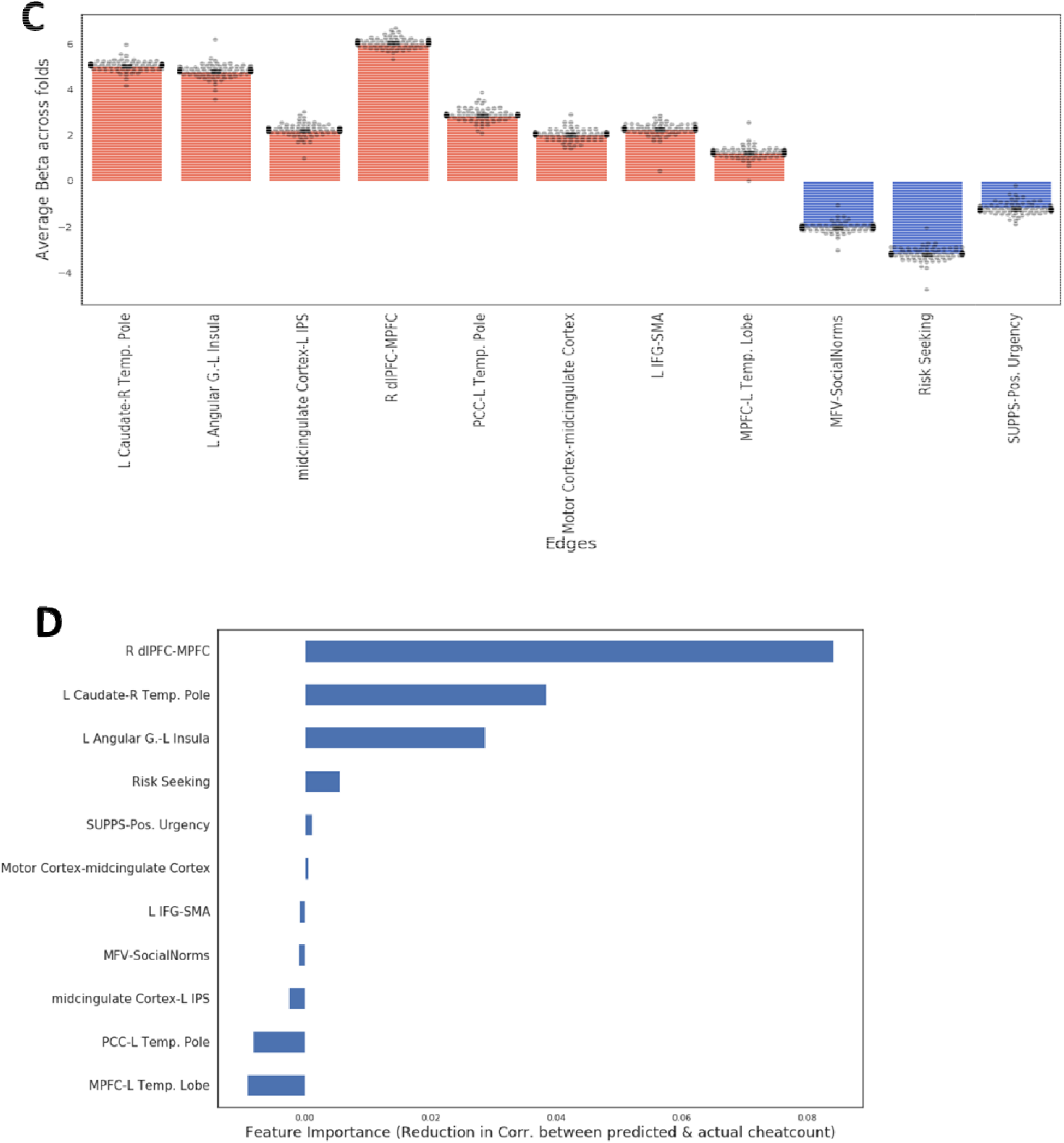
Comparing the contribution of the functional connectome and questionnaires as predictors of honesty. A) Barplot depicting the beta coefficients of the negative predictors from the model predicting honesty, fitted on questionnaires only, averaged across folds, that were significantly different from zero (individual points reflect beta at each fold). B) Correlation between predicted and actual level of honesty for the questionnaire, neural and combined model in comparison. C) Barplot of beta coefficients averaged across folds from the combined model (Neuro + Questionnaires) predicting honesty, that were significantly different from zero (individual points reflect beta at each fold). D) Feature importance as calculated as the reduction in correlation between predicted and actual honesty when removing a given predictor for all predictors of the combined model that were nonzero at each fold.

To see whether a model based only on questionnaires, only on neural data, or a combined model performs best in predicting out-of-sample honesty, we then directly compared these different models. As this analysis is concerned with selecting only the most important predictors among several competing candidates, we used a lasso regression approach. The lasso regression adds a penalty term to the equation which shrinks coefficients in the model to zero and thus reduces complexity of the model and multicollinearity of predictors (Tibshirani, 1996). In this way it also selects the most important predictors in the model. The lasso-CPM on the questionnaire data alone resulted in a significant prediction (*r* = 0.36, *p* < 0.05; see figure 6B). Using lasso-regression on only the neural data using the eight significant connections identified above (see Figure 5C), a significantly higher (as compared to the questionnaire data: *z* = −2.88, *p*<0.005) out of sample prediction was observed (*r* = 0.67, *p* < 0.05; see figure 6B), as compared to the questionnaire data.

Next, we implemented a combined lasso where both the neural (the eight significant predictors from the multiple regression CPM, see Figure 5C) and questionnaire predictors were added to the lasso regression model and we observed a similar out-of-sample prediction score as for the neural-only model (*r* = 0.64, *p* < 0.05). Thus, the neural model clearly outperforms the questionnaire model and adding the questionnaire measures to the neural model to form a combined model does not further improve out-of-sample prediction (*z*=-0.35, *p*=0.35).

To test the contribution more formally, we also used a permutation importance approach (as explained in the method section) to explore which predictors contribute most to the out-of-sample prediction accuracy. This permutation importance analysis revealed that the neural predictors are indeed more important for the predictive performance (correlation between predicted and actual honesty) than the questionnaire data (See Figure 6D). In particular, removing the functional connectivity between the right dlPFC and the MPFC resulted in a reduction of the correlation with 0.08, which is twice as large as any of the other predictors and highlights the importance of this predictor. In addition, the connectivity between the right Temporal Pole and the left Caudate Nucleus, and the connectivity between the Angular Gyrus and the Insula, were important predictors of honesty (see Figure 6D). The reason for some of the edges showing negative permutation importance may be due to multicollinearity between the predictors as can be seen in the Appendix 5 (see Figure 9).

## Discussion

Many of the economically most costly forms of unethical behavior such as tax evasion, stock manipulations or movie and music piracy relate to the moral domain of (dis)honesty, in which unethical behavior is not targeted at a clearly identifiable victim. Large individual differences in (dis)honesty have been observed in this type of behaviour (Gino et al., 2012; 2014, Speer, Smidts & Boksem, 2020), but as of yet, the neural manifestation of this heterogeneity has remained elusive.

Employing connectome based predictive modelling (CPM) in combination with our novel Spot-The-Differences task which exposes participants to the opportunity to cheat, we identified a functional connectome that is predictive of (dis)honesty at the individual level. More precisely, we were able to accurately predict how much an unseen participant would cheat based on her brain’s functional organization. Specifically, we observed a correlation between predicted and actual cheatcount (r = 0.61) that substantially exceed the typical range of correlations (between r = 0.2 and r = 0.5) reported in previous studies employing CPM (Shen et al., 2017). To further test the predictive accuracy of the resting state network we also performed binary classification using several different splits and classifiers to determine whether a participant is a cheater or not and again found high and significant classification accuracies (up to 89%, see Appendix 7). Notably, the predictive accuracy was significantly higher using the brain’s functional connectome than using personality questionnaires.

By combining CPM with (regularized) multiple regression models, we were able to demonstrate that particularly functional connectivity between the MPFC and the right dlPFC was predictive of honesty. This connection was twice as important than any other region in the honesty network. In addition, connectivity between the left caudate and the right temporal pole, the left angular gyrus and the left insula, the midcingulate cortex and the intraparietal sulcus, the PCC and the left temporal pole, the motor cortex and the midcingulate cortex, the left IFG and the SMA, and the MPFC and the left temporal pole contributed significantly to predicting honesty.

In light of previous research on moral decisions, the regions we identified can be associated with three networks frequently found to be involved in moral decisions making. Firstly, the right dlPFC, which has been associated with inhibiting selfish impulses (Speer & Boksem, 2019; Yamagishi et al., 2016; Strang et al., 2014; Steinbeis et al., 2012) and increasing honesty (Marechal, Cohn, Ugazio, & Ruff, 2017), and the left IFG, linked to inhibition of predominant responses (Wager et al., 2005; Verbruggen and Logan, 2008; Sharp et al., 2010; Stokes et al., 2011), can be considered nodes in the cognitive control network. In contrast, the MPFC, the temporal pole, the PCC, the Angular Gyrus and the intraparietal sulcus have consistently been associated with self-referential thinking (Gusnard et al, 2001; Meffert et al., 2013; Van Buuren et al., 2010). This self-referential thinking network has previously been found to be more strongly activated when honest people are exposed to an opportunity to cheat and more strongly interconnected when honest people make an honest decision (Speer, Smidts & Boksem, 2020). Lastly, the caudate nucleus, which has been found to be involved in anticipation and valuation of rewards (Ballard & Knutson, 2009; Knutson, Adams, Fong, & Hommer, 2001; Abe & Greene, 2014) can be considered important nodes in the reward network (Bartra et al., 2013). Participants with higher levels of activation in the reward network, in anticipation of rewards, have indeed been found to be more dishonest (Abe & Greene, 2014; Speer, Smidts & Boksem, 2020).

To confirm that the identified regions indeed belong to the networks proposed, we conducted a conjunction analysis between our results and meta-analytically derived maps associated with self-referential thinking, reward, and cognitive control, obtained using Neuroquery (Dockes et al., 2020). The conjunction analysis revealed that there is substantial neural overlap between our results and the meta-analytically derived maps (see Appendix 6), which validates our interpretation of the observed functional networks and reduces the reverse inference problem (Poldrack, 2006).

Our findings suggest how neurocognitive processes shape individual differences in moral choices. Specifically, we found that honest participants exhibited higher connectivity within the self-referential thinking network, but also between self-referential thinking and cognitive control and reward regions. Firstly, these findings suggest that the self-referential thinking network may represent the moral self-concept and may be engaged in self-concept maintenance when exposed to an opportunity to cheat (Mazar, Amir & Ariely, 2008). Secondly, our results suggest that increased resting state functional connectivity between self-referential thinking and cognitive control regions may enhance cognitive control processes to facilitate achieving the long-term goal of maintaining a positive self-concept. Lastly, stronger connectivity at rest between self-referential thinking regions and reward regions may reflect integration of the appeal of maintaining a positive self-concept into a representation of value.

Collectively, our findings suggest that a person’s moral choices depend on how well the self-referential thinking network is functionally connected to cognitive control and reward networks. Stated differently, stronger interaction between our self-concept, cognitive control and reward processes may be essential for honest behavior. While these inferences need to remain speculative, as we can not determine the directionality of these functional connections, these findings align well with self-concept maintenance theory (Mazar et al., 2008), as they suggest that self-referential thinking is the main driving force for honesty.

We did not find evidence supporting our hypothesis that stronger connectivity within the reward network or increased negative coupling between the reward and cognitive control network are predictive of dishonesty. One explanation for the failure to confirm our hypothesis may be that the extent to which we anticipate rewards and incorporate reward processes into our (moral) decisions may be similar across individuals, whereas it is the extent to which our moral self-concept is developed and the degree to which self-referential thinking processes, in concert with cognitive control processes, guide our moral decisions, may vary substantially between individuals thus driving heterogeneity in (dis)honesty.

We also found that several well-established self-report personality measures of impulsivity correlated significantly with dishonesty on our task. This highlights the ecological validity of the Spot-The-Difference task as a measure of dishonesty, as impulsivity has frequently been associated with other forms of cheating such as academic cheating (Cochran et al., 1998; Anderman et al., 2009), dishonesty more generally (Gino et al., 2011) and unethical behavior (Zimmerman, 2010; Loeber et al., 2014). While we found that these self-reported measures of impulsivity were significantly correlated with dishonesty, the predictive power of these measures was outperformed by the predictive accuracy of the functional connectome. In a direct comparison, the model including only neural measures achieved a correlation between predicted and observed rate of dishonesty that was nearly twice as high as the model containing self-report measures only.

In addition, a permutation importance analysis (Breiman, 2001) revealed that in a model combining self-report and neural measures, mainly the neural measures contributed to the predictive accuracy. This dominance of neural measures may be in part by due to the fact that self-report measures may suffer from social desirability bias particularly in the context of dishonesty (Grimm, 2010). Participants may not want to admit that they are impulsive or may not even be aware that they are. This social desirability bias might also explain why we unexpectedly observed that participants who report to be more sensitive to social norms and matters of purity also cheat more. Some participants, particularly the dishonest ones, might have overstated how much they care about, and base their decisions on, social norms and moral purity.

The superiority of the neural measures might be attributed to the fact that they are uncontaminated by the biases above and may represent neuro-cognitive processes that govern our behavior in a way that we often may not be aware of. An individual’s functional connectome has been shown to be largely task-independent, robust and reproducible across time (Finn et al., 2015). Thus, when combining CPM on rsfMRI data with incentivized tasks simulating real world behavior, the functional connectome may be predictive of a range of traits and behavioral tendencies. For instance, recent research using CPM has used heterogeneity in functional connectivity to predict measures such as intelligence (a well-established and reliable cognitive trait, Finn et al., 2015), creativity (Beaty et al., 2018), as measured by the Alternative Uses task (AUT, Guilford, 1967; a well-established creativity task evoking actual creative behavior), impulsivity in an incentivized delay-discounting task (Li et al., 2013), and trust behavior in an incentivized trust game (Hahn et al., 2014). The functional connectome at rest, when combined with a reliable and valid measure of behavior, seems to provide direct and reliable access to different behavioral tendencies that is not distorted by response biases questionnaires are susceptible to. Importantly, it permits measuring the neural manifestation of socially inadmissible behavioral tendencies such as dishonesty, that could be concealed in self-report measures. Here, we again demonstrated that this approach is particularly useful when studying complex and socially inadmissible behaviors such as dishonesty that are not limited to a specific brain region but require sophisticated interaction between a distributed network of regions across the brain. Therefore, it can be concluded that resting state functional connectivity combined with CPM is a highly useful tool for studying individual differences, specifically when these differences pertain to socially undesirable behavior.

Importantly, due to the fact that CPM is based on cross-validation, it is in two ways superior to previous correlational research investigating the relationship between functional connectivity or personality tests and human behavior: Firstly, from the perspective of scientific rigor, cross-validation is a more conservative approach to infer the presence of a relationship between brain and behavior than is correlation, due to the fact that it is designed to prevent overfitting by means of testing the strength of the relationship in a novel sample. It thus increases the replicability of findings in future studies. Secondly, regarding the practical perspective, establishing predictive power is crucial to translate neuroimaging insights into tools with practical relevance (Shen et al., 2017). Testing performance of models in independent samples, in our case individuals, facilitates the evaluation of the generalizability of findings and consequently the eventual development of neuroimaging-based biomarkers with real-world utility. The practical relevance of combining CPM with resting state fMRI is further strengthened by the fact that the acquisition of a resting state scan only requires 7-8 minutes and has been shown to be reliable and reproducible across time (Cao et al., 2014; Zuo and Xing, 2014; Finn et al., 2015), which suggests that only one acquisition is necessary. This reduces the financial cost and administrative complications considerably as compared to task-based neural measures and emphasizes the potential practical relevance as a biomarker for dishonesty that may be a significant first step in developing more sensitive interventions to reduce cheating.

(Dis)honesty remains a complex behavior that requires further research to reveal its many manifestations in the brain. Although honesty is a central moral principle, it is just one of several moral foundations (in addition to Honesty/Cheating, other moral foundations relate to Harm/Care, Loyalty/Betrayal, Authority/Subversion, Sanctity/Degradation; Graham et al., 2011). Therefore, an interesting avenue of research would be to explore whether resting state functional connectivity can predict unethical behavior in several different domains of morality. More specifically, it could be investigated whether heterogeneity in the same functional connectomes similarly predict (dis)honesty, harm aversion or loyalty, or whether personal tendencies regarding these distinct but related moral domains are represented by different functional networks in the brain.

In summary, our extended CPM model applied to a large and diverse sample revealed that self-referential thinking processes in interaction with cognitive control and reward processes are of central importance in promoting honesty. We showed that an individual’s propensity to cheat depends on how well the self-referential thinking network is connected to cognitive control and reward networks at rest. These individual differences in resting state connectivity between self-referential thinking, reward and cognitive control network can be used to reliably and accurately predict people’s tendency to cheat and substantially exceed the predictive accuracy of self-report measures.

## Appendices

## Appendix 1: Visual search task

To further increase the credibility of our cover story on brain processes underlying visual search, we also included the visual search task introduced by Treisman and Gelade (1980) at the beginning of our experiment. Specifically, participants were told that the experiment would start with a simple visual task and then proceed to visual searches in more complex visual stimuli in the second task. In this first task, the goal was to determine whether a specific target was present or absent. In each trial participants were presented with colored letters presented in random locations on the screen. If the target was present, then participants had to press the left mouse button as quickly as possible. If no target was present, then they had to press the right mouse button as quickly as possible. For this task, participants had to search for a green T. Participants were instructed to answer as quickly as possible while still being as accurate as possible. The task took approximately 5 minutes and was not analysed as it was included solely for the purpose of increasing the credibility of our cover story.

**Figure 7.**
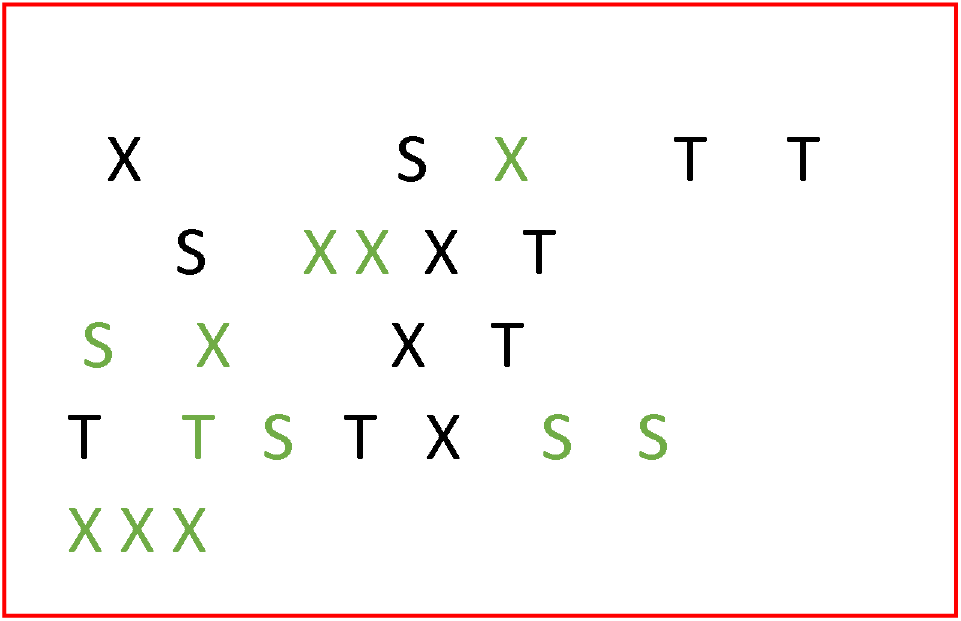
One trial of the simple visual search task. Participants have to indicate whether a green T is among the letters on the screen.

## Appendix 2: Validation of the picture set

Stimuli for the task consisted of 144 Spot-The-Difference image pairs that were downloaded from the Internet. Cartoon images of landscapes containing several objects were selected, to make them engaging and challenging enough for the participants. Landscapes were chosen as they generally satisfied the necessary criteria of containing several different objects, which made the task of spotting differences more challenging and engaging. The stimuli consist of pairs of images that are identical apart from a certain number (1-3) of differences that were created by the experimenter using Adobe Photoshop. Differences consisted of objects added to or removed from the landscape picture or changed colors of objects.

To make sure that participants would be able to find the differences between the images in a reasonable amount of time, we ran a pilot study on Amazon’s Mechanical Turk with 205 subjects using 180 pictures to test the difficulty to spot the differences between the images and to determine the optimal duration of picture presentation. Participants were presented with cartoon image pairs, presented horizontally next to each other, containing three differences and were asked to click on the differences identified in the image on the right hand side. They were given 15 seconds to make their response. Using the heatmap function provided by Qualtrics, regions of interest were defined around the locations of the differences in the image on the right hand side and response times for each of the clicks were recorded. This allowed us to test whether participants were able to find all differences in an image pair, which differences were particularly difficult to find, and how long it took to identify all differences. Based on the responses of these 205 participants, 36 image pairs that took too long or had differences that were too difficult or too easy, were removed, resulting in 144 images that took 92% participants less than 6s to find all three differences (M=5.4s, SD =1.5s).

## Appendix 3: Connectome Based Predictive Multiple Regression without suspicious participants

To test for the robustness of the CPM model and potential effects of suspicion on the reported effects we conducted the same analysis as above on the data of participants who were not suspicious of the purpose of the study. The analysis revealed that using the CPM multiple regression approach as above we are still able to significantly predict cheatcount from the negative connectome (functional connection that are negatively correlated with cheating) with high accuracy (*r* = 0.39, *p* < 0.05). As before no significant prediction was possible for the positive connectome. These findings demonstrate that there does not seem to be a significant effect of suspicion about the purpose of the task on the reported effects and highlight the robustness of our findings.

## Appendix 4: Correlations between personality scores and cheatcount

**Figure 8.**
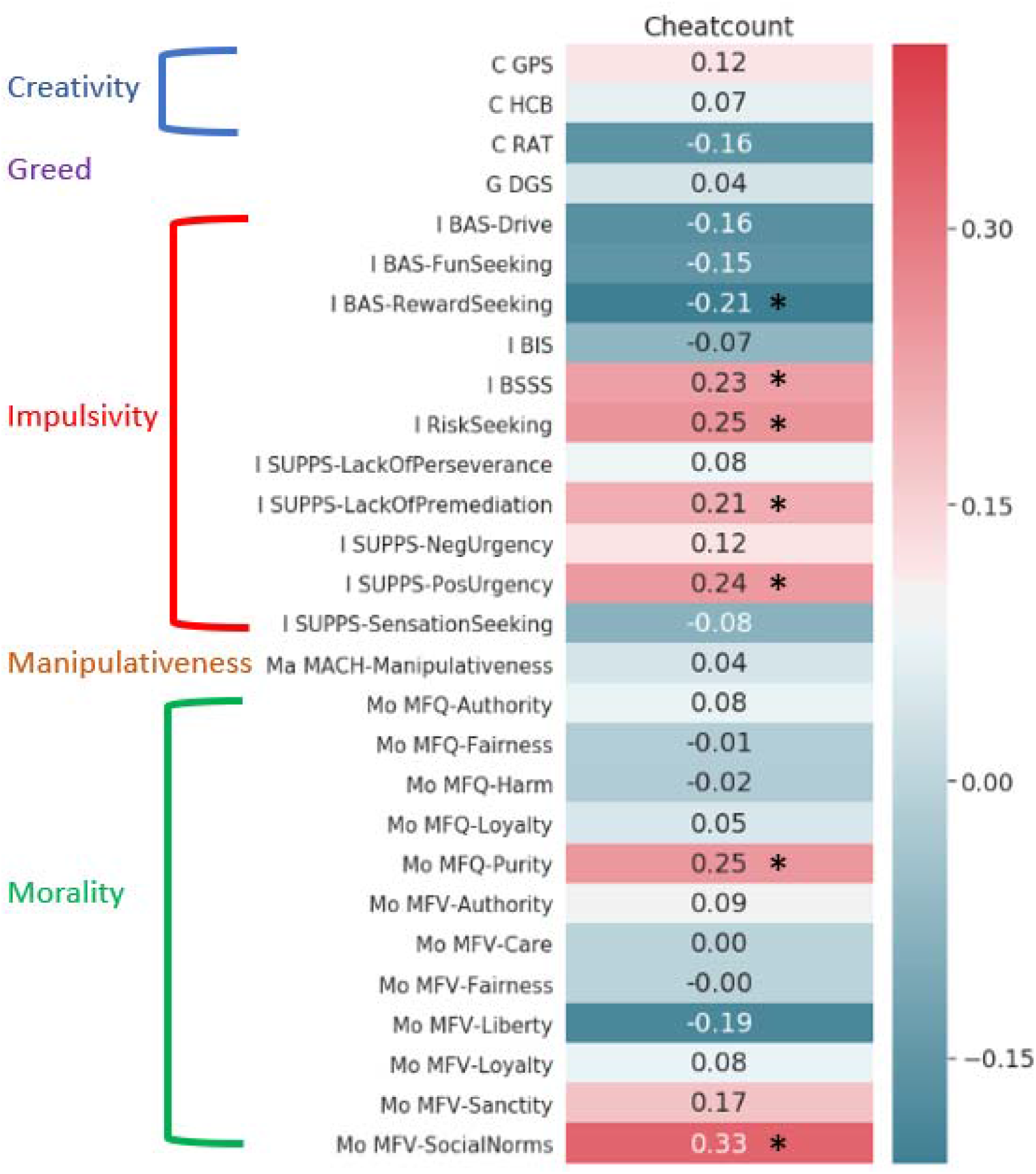
Correlations between all personality measures used and the cheatcount. C = Creativity; I = Impulsivity; Ma = Manipulativeness; Mo = morality; GPS= Gough’s Personality Scale; HCB = Hovecar’s Creative Behavior; DGS = Dispositional Greed Scale; BAS = Behavioral Approach System; BIS = Behavioral Inhibition System; BSSS= Brief Sensation Seeking Scale; SUPPS = Short Urgency, Premediation (lack of),Perseverance (lack of), Sensation Seeking, Positive Urgency Impulsive Behavior Scale; MACH = MACH-IV test of Machiavellianism; MFQ = Moral Foundations Questionnaire; MFV = Moral Foundations Vignettes.* = indicates significant correlations (p<0.05 uncorrected).

## Appendix 5: Multicollinearity of the Combined CPM lasso model

In order to test why the Questionnaire data does not add additional predictive power to the model when combined with the neural predictors, we computed the correlations between all nonzero predictors of the combined CPM lasso model. As depicted in the graph below (see Figure 9) there is a significant negative correlation between the sensitivity to social norms (MFV-SocialNorms) and the functional connectivity between the Angular Gyrus and the left Insula (*r* = −0.33, *p* < 0.05), the right dlPFC and the MPFC (*r* = −0.25, p < 0.05) and the PCC and the left Temporal Pole (*r* = −0.24, p < 0.05). In addition, significant correlation was found between risk seeking and the functional connectivity between the Motor cortex and the midcingulate Cortex (*r* = −0.29, *p* < 0.05). Lastly, significant correlations were found between the Positive Urgency subscale of the S-UPPS scale and functional connectivity between the midcingulate Cortex and intraparietal sulcus (IPS; *r* = −0.27, *p* < 0.05), the PCC and the left Temporal Pole (*r* = −0.21, *p* < 0.05) and the left IFG and SMA (*r* = −0.24, *p* < 0.05). These correlations may explain why adding the questionnaires data did not improve the model as the variance explained by even the most important questionnaire measures (MFV-Social Norms, Risk Seeking, S-UPPS Positive Urgency) was already accounted for by the neural predictors.

**Figure 9.**
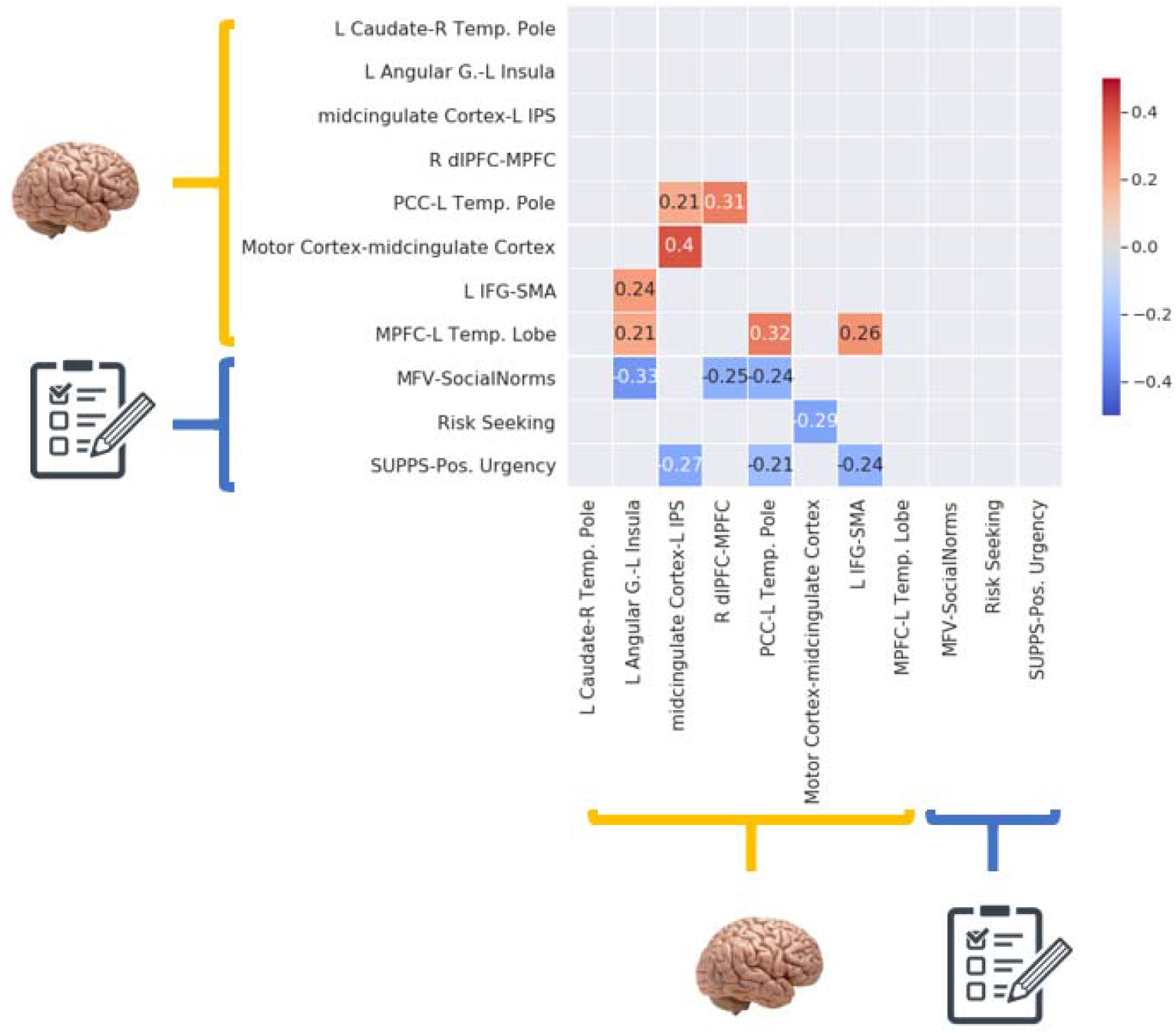
Correlation between predictors in the combined CPM lasso model.

## Appendix 6: Conjunction Analysis to reduce reverse inference

In order to reduce the reverse inference problem (Poldrack, 2006), we also assessed the neural overlap between the regions identified in our analysis and meta-analytically derived maps associated with, respectively, self-referential thinking, cognitive control obtained using Neuroquery (Dockes et al., 2020). Neuroquery is a new meta-analytic tool for human brain mapping that was developed by researchers who were also involved in creating Neurosynth (Yarkoni et al., 2011). The advantage of Neuroquery is that it is focused on producing a brain map that predicts where in the brain a study on a particular cognitive process is likely to report observations, while Neurosynth tests the consistency of observations reported in the literature. Prediction, as opposed to statistical testing, is important because it can be applied *out of sample* and is thus more generalizable. The conjunction analysis reveals that there is indeed overlap between our results and the meta-analytically derived maps.

**Figure 11.**
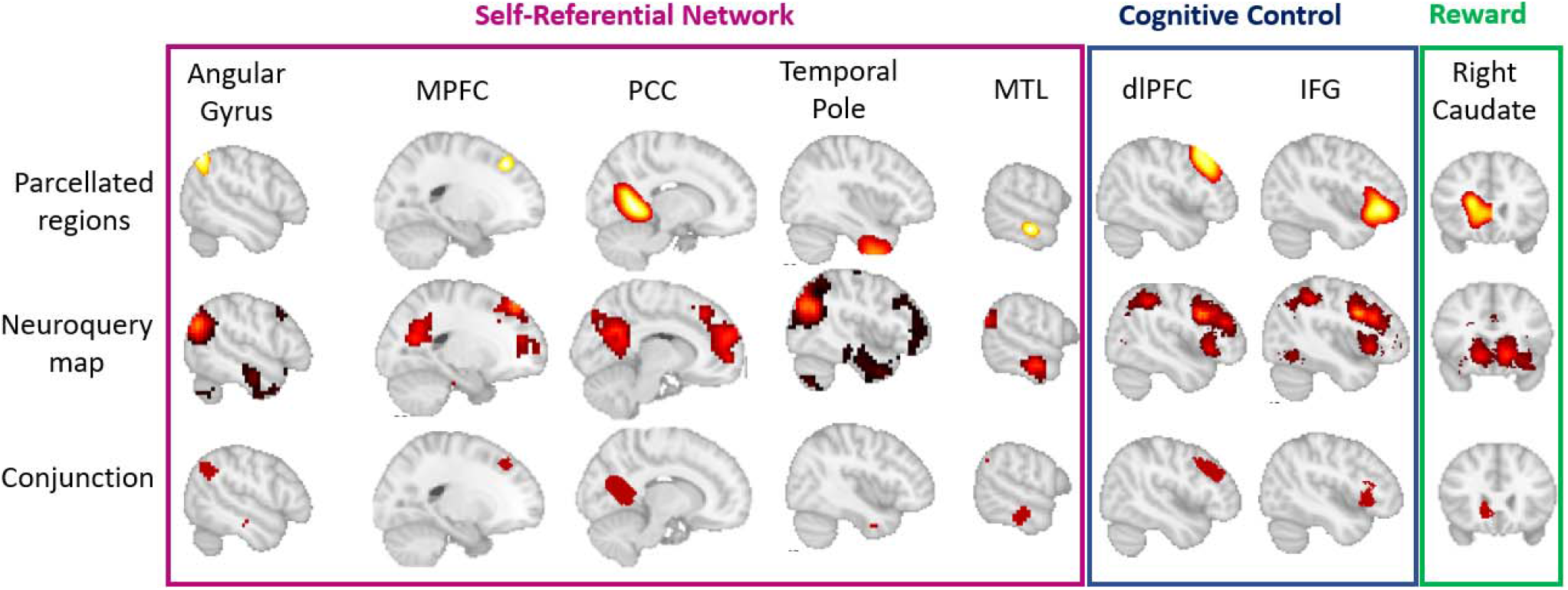
Neural overlap (bottom row) between the regions identified to be important in predicting (dis)honesty (top row) and meta analytically derived maps for self-referential thinking cognitive control and reward processing (middle row).

## Appendix 7: Binary Classification Analysis using the top 8 connections

In order to get a better understanding of the predictive accuracy of the predictors identified in the multiple regression CPM we conducted a binary classification using the 8 most predictive connections. To this end, we first performed several types of splits on the participants’ cheatcount: A median split (Median = 20), a 3-way split keeping the lowest 33% and the highest 33% (33% = 9, 66% = 35.36), a 4-way split keeping the lowest quartile and the highest quartile (25% = 7, 75% = 44), and a 5 −way split keeping the lowest two quintiles and the highest two quintiles (20% = 5.6, 40% = 14.2, 60% = 28, 80% = 52.2). The reasoning behind this approach was to see how well we can distinguish honest participants from cheaters while trading off between sample size (highest using the median split) and discriminability (highest in the 4-way split). More precisely, the discriminability is lowest in the median split because all participants that are close to the median and are hardest to classify are kept, whereas in the 4-way split only the most extreme participants are kept. Subsequently, different classifiers commonly used for binary classification, including a logistic regression classifier, a random forest classifier (Breiman, 2001) and a support vector classifier (Cox & Savoy, 2003) were trained on the functional connectivity patterns of each participant to determine whether a participant was a cheater or an honest participant. In order to avoid overfitting and inflated prediction accuracy (Vul, Harris, Winkielman, & Pashler, 2009) this was done using leave-one-out cross-validation as in the CPM analysis in the main text. Significance was estimated using permutation testing (N=1000). The classification analysis revealed that we could significantly classify an unseen participant as a cheater or an honest individual with a classification accuracy ranging between 71% for an SVM trained on median split data to a lasso regression model trained on 4-way split data with a classification accuracy of 89% (see Table 1).

**Table 1.** Classification scores (%) across different splits and classifiers.

|  |  |  |  |  |
| --- | --- | --- | --- | --- |
|  | **SVM** | **Lasso** | **RandomForest** | **N** |
| **Median** | 71 | 77 | 76 | 99 |
| **3-way** | 80 | 80 | 77 | 65 |
| **4-way** | 85 | 89 | 81 | 48 |
| **5-way** | 80 | 84 | 79 | 79 |

SVM = support vector machine, N = number of subjects left after the split. All classifications were significant at $p < 0.01$ estimated using permutation testing (N=1000).

